# Emergence of ribozyme and tRNA-like structures from mineral-rich muddy pools on prebiotic earth

**DOI:** 10.1101/2020.02.12.944926

**Authors:** Suvam Roy, Niraja V. Bapat, Julien Derr, Sudha Rajamani, Supratim Sengupta

## Abstract

The RNA world hypothesis, although a viable one regarding the origin of life on earth, has so far failed to provide a compelling explanation for the synthesis of RNA molecules with catalytic functions, from free nucleotides via abiotic processes. To tackle this long-standing problem, we develop a realistic model for the onset of the RNA world, using experimentally determined rates for polymerization reactions. We start with minimal assumptions about the initial state that only requires the presence of short oligomers or just free nucleotides and consider the effects of environmental cycling by dividing a day into a dry, semi-wet and wet phases that are distinguished by the nature of reactions they support. Long polymers, with maximum lengths sometimes exceeding 100 nucleotides, spontaneously emerge due to a combination of non-enzymatic, non-templated polymer extension and template-directed primer extension processes. The former helps in increasing the lengths of RNA strands, whereas the later helps in producing complementary copies of the strands. Strands also undergo hydrolysis in a structure-dependent manner that favour breaking of bonds connecting unpaired nucleotides. We identify the most favourable conditions needed for the emergence of ribozyme and tRNA-like structures and double stranded RNA molecules, classify all RNA strands on the basis of their secondary structures and determine their abundance in the population. Our results indicate that under suitable environmental conditions, non-enzymatic processes would have been sufficient to lead to the emergence of a variety of ribozyme-like molecules with complex secondary structures and potential catalytic functions.

## 1 Introduction

RNA polymers possess the ability to replicate and store information like DNA. These characteristics together with the discovery of ribozymes [1–3] and regulatory RNA like riboswitches [4–8] that responds to changes in concentrations of a variety of small molecules provide indirect evidence of an RNA world that preceded life based on DNA and proteins. Despite such promising indirect evidence, several major challenges [9–13] remain for the RNA world to be considered as a realistic epoch. The functional RNA molecules present in the ribosome as well as the non-coding RNA regulators are all synthesized with the help of enzymes within a living cell. Even though several ribozymes have been synthesized in the lab primarily using *in vitro* selection experiments [14, 15], a major challenge of the RNA world hypothesis lies in demonstrating the ability to abiotically synthesize not just the RNA polymers but also their monomer building blocks through non-enzymatic processes. Recent progress on abiotic synthesis of pyrimidine ribonucleotides [16–19] has provided deep insights into the chemical origins of such informational molecules. Nevertheless, much work remains to be done for a complete understanding of the emergence of functional biomolecules via prebiotic synthesis. In addition to the challenges of synthesis, the evolution of such biomolecules on primordial earth also has to circumvent the error-threshold problem [20] to ensure that they are not subject to mutational degradation.

The non-enzymatic synthesis of long biopolymers depends crucially on environmental conditions. There has been much speculation on ideal prebiotic environments that can be favourable for emergence of life that either invoke terrestrial geothermal pools [21] or hydro-thermal vents present on the ocean floor [22, 23]. While a purely aqueous environment promotes diffusion useful for enhancing monomer availability required for polymer extension, it nevertheless leads to the hydrolysis of the existing polymers. Additionally, it results in dilution of the system making collisions required for polymerization less favourable. On the other hand, polymerization reactions are thermodynamically favourable under dry conditions [24] that also reduce breaking up of existing polymers by hydrolysis of phosphodiester bonds. But continuous dryness reduces the diffusivity of the molecules and hence reduces the reaction rates. Significant increase in the likelihood of formation of RNA-like polymers is observed in the presence of alternate wet-dry cycling conditions [25–30] prevalent in a primordial earth. In addition to concentrating the starting monomers, the dehydration phase also enhances loss of water, thus promoting bond formation by facilitating the condensation reaction. This is thermodynamically unfavourable at room temperature in bulk water. The subsequent rehydration phase also facilitates the re-distribution of the monomers and oligomers, consequently increasing the overall efficiency of the reaction. Significantly, when lipids are involved in such wet-dry cycling processes, they also protect the resultant RNA oligomers from degradation. Thus, such conditions have been shown to be favourable for the formation of long polymers (25-100 nucleotides) especially in the presence of lipids [25–27] and salts [31, 32] like *NH*_4_*Cl*. Montmorillonite clay has also been suggested as a substrate [33, 34] that can enhance polymerization rates resulting in synthesis of 50-mer long polymers. Nevertheless, non-templated polymer extension rates achieved on clay are still a couple of orders of magnitude less than template-directed polymer extension rates. Such non-enzymatic template-directed primer extension rates have been measured under various circumstances [35–42] albeit with activated nucleotides, and suggested as a viable alternative mechanism for synthesizing long polymers.

Due to the time-limiting nature, relatively low yields of most non-enzymatic reactions and limitation of our knowledge of prebiotic environmental conditions, experimental investigations are often limited by the questions they can plausibly address. Considerable insights into prebiotic evolutionary processes can be derived by supplementing experimental work with theoretical modeling. It is therefore not surprising that several theoretical models have been proposed to understand the origins of functional RNA polymers. Obermayer et al (2011) showed that complex RNA replicators can emerge in an RNA reactor with a thermal gradient that allows for spontaneous ligation of RNA strands [43]. Thermal gradient helps in accumulation of monomers, which in turn helps the formation of longer RNA strands, thereby increasing their abundance in the population [44, 45]. Wet-dry cycles also facilitate formation of long polymers [46]. Monomer availability can control the dynamics of the system in the sense that lower rate of monomer flow can favor the dominance of more complex sequences [47]. Derr et al (2012) showed that with a limited number of monomers, template-directed ligation leads to a diverse pool of RNA strands, where the individual strands also have high compositional diversity [48]. Walker et al (2012) analysed the role of cycling and diffusion in a model of prebiotic polymer formation through template-directed process evolving in a flat fitness landscape [49]. Higher rates of template-directed ligation were shown to lead to emergence of sequences having the same type of chirality [50]. Most theoretical models however rely on deriving conclusions through exploration of parameter space that is unconstrained by experiments.

Our main aim in this paper is to use recent experimental data on non-enzymatic rates of polymer extension with and without templates to develop and analyse a realistic computational model of prebiotic polymer formation. In the process, we explore the conditions under which long informational polymers, with the kind of structural complexity observed in ribozymes, can emerge. The chemical processes we examine would have occurred during the earliest epoch of the RNA world and eventually led to the emergence of RNA replicases that were instrumental in accurate replication of strands. The latter epoch had to contend with competition between selfish (those ribozymes that cannot replicate themselves and depend on other altruistic replicases for their synthesis) and altruistic RNA replicases (those ribozymes that can replicate themselves as well as their selfish counterparts) and uncover protocols, either through spatial clustering or through confinement in protocells, that prevent unrestricted proliferation of selfish replicases [51–61]. We allow for replication of RNA molecules via a template-directed primer extension process in which an RNA strand can act as template to extend a small primer by attaching monomers to its leading end in a manner that results in complementary base pairing with the corresponding nucleotides of the template. Upon full extension the primer sequence will be complementary to the template. But such processes are prone to errors, where at a low rate, non-complementary monomers can also attach to the primer resulting in a higher mutation rate rate in such systems [36]. Additionally, this effect has also been shown to increase in the presence of prebiotically pertinent co-solutes [35]. We examine the consequences of reduced rates of primer extension after a mismatch [41] on the length and type of sequences generated by using primer extension rates from experiments [35, 62]. We also incorporate the relatively slower but nonetheless vital non-templated monomer/polymer extension process by addition of a single monomer (concatenation) and show that both these processes are essential for generating long, plausibly functional polymers with ribozyme-like structures.

To mimic the effect of prebiotic environmental cycles on polymer extension, we divide each day into different phases based on temperature and dryness, where different reactions require different optimum temperatures for their occurrence. Even though the average length of a day on the early Earth is thought to have been shorter than today [63], our choice of 24 hours for the duration of a day lies within the bounds of reason. The dry phase is preferable for spontaneous concatenation of polymers [Figure-1(A)] whereas the wet phase promotes hydrolysis of existing polymers. We first present a 2-phase analytical model, well-supported by numerical simulations, to show how the effect of cycling between wet and dry phases can lead to increase in average length of the synthesized polymers. Template-directed concatenation processes [Figure-1(B)] are promoted under environmental conditions that allow for a supply of monomers to extend the primer attached to the template while at the same time suppressing the breaking of templates due to hydrolysis. Such conditions are prevalent in a semi-wet phase which motivated us to introduce a more realistic 3-phase model where such a semi-wet phase is sandwiched between the dry and wet phases. We also incorporate structure-dependent hydrolysis of existing sequences by imposing different hydrolysis rates for the bonds that link unpaired nucleotides relative to the bonds that link singly or doubly-paired nucleotides (see Figure-2) in the 3-phase model. We keep track of the structural diversity of the RNA sequences during the entire simulation.

**Figure 1:**
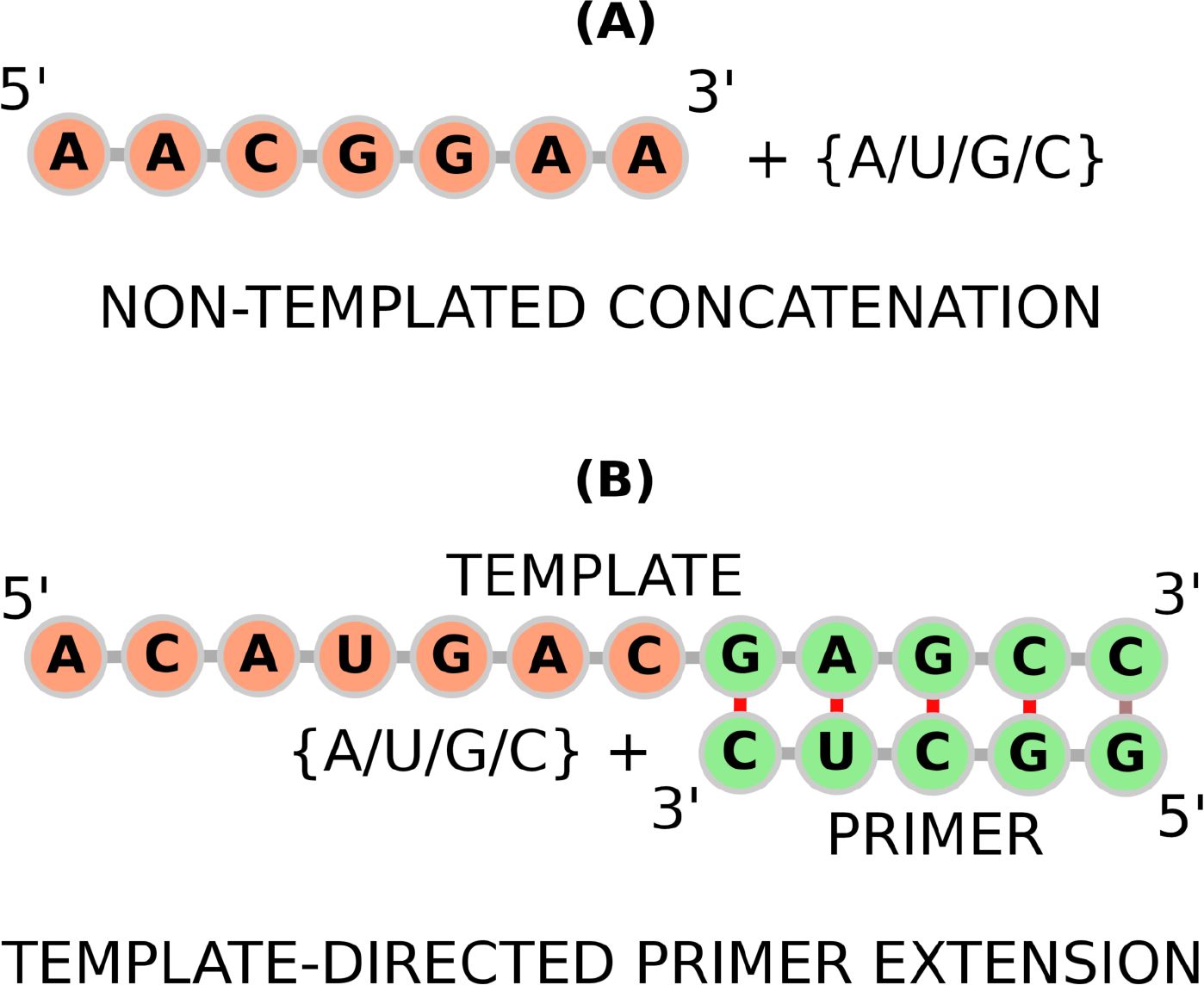
Pictorial representation of **(A)** non-templated concatenation process and **(B)** template-directed primer extension process

**Figure 2:**
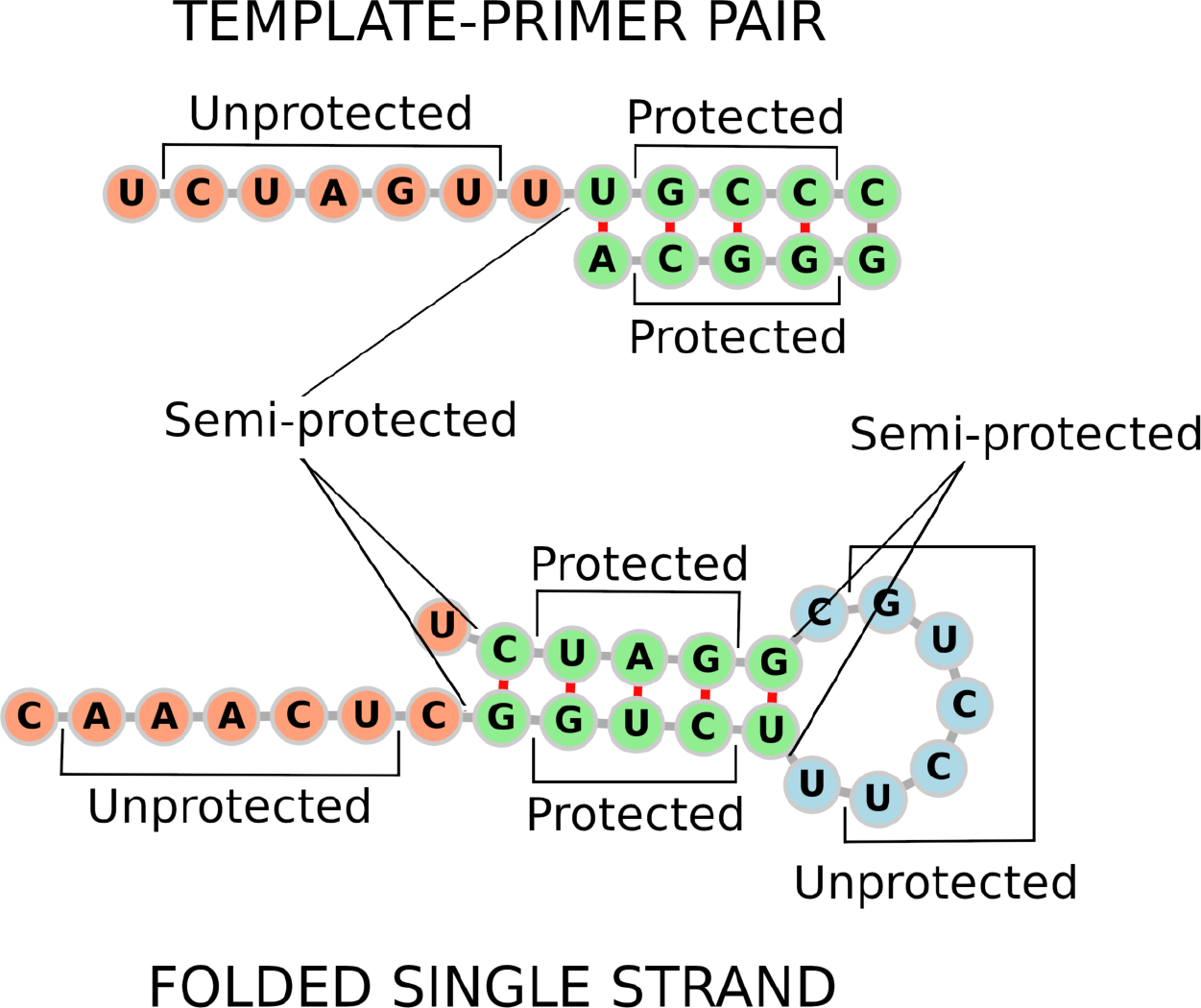
Different types of phosphodiester bonds classified by their propensity to be hydrolysed, depending on the presence of paired or unpaired nucleotides on either side of the bond. If both nucleotides are unpaired, the hydrolysis rate is *p*_*uu*_ *K*^*hyd*^; where as if both of them are paired, the hydrolysis rate becomes *p*_*pp*_ *K*^*hyd*^. If any one of the neighboring nucleotides is paired, then hydrolysis occurs at a rate *p*_*up*_ *K*^*hyd*^.

Our simulations generated moderately long RNA strands with average length ~ 40 nucleotides, with a small fraction of sequences extending even beyond 100 nucleotides, under certain conditions. To better understand sequence diversity, we classified the sequences on the basis of their secondary structures. We found that longer sequences (>20mers) often possessed complex foldable structures (single hairpin, double hairpin, hammerhead and cloverleaf) of the type observed in ribozymes and tRNA. Under certain conditions we also obtained long replicated copies (maximum length >30 nucleotides) of strands by the template-directed primer extension process which indicate how double-stranded RNA molecules can also emerge via such process. Our work provides conclusive evidence that, subject to certain environmental constraints, long RNA polymers may have been readily produced due to a combination of both slower non-templated concatenation reactions and relatively faster template-directed primer extension processes. Many of the generated sequences were found to possess complex structures that may plausibly be recognized as a proxy for their catalytic capabilities.

## 2 Methods

We choose the location of our system to be a muddy area in the prebiotic earth (as found in the regions around geothermal pools), which undergoes periodic hydration and dehydration due to the effect of day-night cycles. Experiments typically use *O*(10 *mM*) monomer concentrations and *O*(1 *μM*) polymer concentrations, which means a 1 *μm*^3^ volume has *O*(10^6^) monomers and *O*(10^2^) polymers on an average. Hence we consider a small open volume of size ~ 1 *μm*^3^ as our system, which describes a tiny fraction of macroscopic volumes, but sufficient enough to extract the
system level properties from it. In the next section, we study a scenario where each day is divided into 2 phases, a hot, dry phase followed by a cold, wet phase. In subsequent sections, we introduce a semi-wet (or semi-cold) phase between a dry (or hot) phase and a wet (or cold) phase. The dry phase has the maximum temperature ~ 90^°^ *C*, (a plausible temperature on primordial earth) and very low amounts of water. Because of it’s high temperature, the dry phase facilitates spontaneous concatenation, where a monomer can concatenate to the 3’ end of another monomer or an existing polymer and form a phosphodiester bond. The system is assumed to be rich in minerals. The concatenation rate in presence of minerals is taken to be consistent with rates reported in experiments [64, 65]. The semi-wet phase sandwiched between the dry and wet phases is characterized by intermediate temperatures intermediate amounts of water. The semi-wet phase is suitable for template-directed primer extension as the formation of hydrogen bonds between the templates and primers requires a lower temperature, which is dependent on the melting temperature (*T*_*m*_) of the primer-template pair in question. The presence of minerals increases the primer extension rates by 3-4 fold [62]. The wet phase is the coldest one in which the system is heavily hydrated by water. The wet phase facilitates hydrolysis with rates that depend on the pH of the medium [66] and also allows for length-dependent diffusion of polymers [67] that preferentially filters out shorter polymers from the system. We assume a slightly alkaline medium (pH = 8). The dry and semi-wet phase therefore promote enhancement of polymer length in contrast to the wet phase which leads to fragmentation of existing polymers. We assume the monomers to be highly diffusive in all 3-phases as they are the lightest molecules in the system. Because of the diffusive nature of the monomers there is a constant inflow and outflow of monomers into and out the system volume, making the monomer concentration effectively constant. Abiotic synthesis of new monomers with time also helps ensure that the monomer concentration remains constant. Hence the concatenation and primer extension rates are independent of the monomer concentrations i.e. these reactions are pseudo 1st order reactions. But unlike the monomers, the diffusion of the polymers is affected by the presence of water in the system, as they are much bigger compared to the monomers. Polymers have highest diffusivity in the wet phase and lowest diffusivity in the dry phase. Hence we neglect non-templated ligation reaction between the polymers in dry phase because of the low diffusivity of the polymers in dry phase and also because of the fact that ligation reaction in general is significantly slower than concatenation reaction (*K*^*con*^ ~ 10^3^ *K*^*lig*^) [64, 68]. Figure-1 depicts the non-templated concatenation and template-directed primer extension processes respectively.

For the 2-phase model described in the next section, we begin our simulations at the beginning of the dry phase while for the 3-phase model (described in subsequent sections), our simulation begins at the onset of a semi-wet phase. We start with 100 homogeneously stacked sequences of lengths normally distributed around 8 nucleotides. Stacked sequences are not formed by concatenation reactions but by condensation of monomers when there are large number of monomers present in the system. The initial strands are homogeneously stacked A or G polymers, in view of the fact that purines can spontaneously aggregate to form poly-A or poly-G sequences of small lengths. We neglect further stacking of monomers during the simulation. Even though we used short stacked sequences and free monomers as initial condition for our 3-phase model, we have verified that the initial stacked poly-A and poly-G sequences have no effect on the final results. All our results also hold for the case when we start the simulations with only free monomers at the beginning of a dry phase. At the beginning of the semi-wet phase we first check the templating efficiency of each strand. A strand of length 10 nucleotides or more, will act as a template with a probability 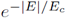, where *E*_*c*_ is critical free energy for folding of the strand into secondary structures. This functional choice of templating efficiency is dictated by the fact that sequences with lower free energy have better folding capability, whereas those with higher free energy are less-likely to fold and therefore act as better templates [69]. The critical free energy for folding into secondary structures is taken as *E*_*c*_ = 2 kcal/mol (as the thermal energy KT ~ 0.6 kcal/mol at normal temperatures and the probability of forming the secondary structures with minimum free energy is usually ~ 0.4 (as found from the ViennaRNA package [70]). Hence we choose *E*_*c*_ > 0.6/0.4 kcal/mol). We choose a set of templates randomly from the set of strands according to their templating efficiencies. We then attach a monomer across the 3’ end nucleotide of each of the chosen templates, to act as their primers, i.e. we begin with primers of length 1 nucleotide only. The type of the attached monomer depends on the nucleotide at the 3’ end of the template, across which it binds and the corresponding relative reaction propensities for addition of the four types of monomers [35]. The primers are then allowed to extend along the 3’ - 5’ direction of the templates by step-wise addition of monomers according to the rates determined from [35, 41]. It is further observed that when there is a mis-incorporation in the previous step, i.e. a non-complementary monomer gets added across the previous templating base, the extension rates are reduced significantly and the probability of another mis-incorporation is increased [41]. We use these experimentally determined rates in our model. Template-directed primer extension rates after correct and incorrect base pairings are given in Table-1 and Table-2 respectively. All of these rates are increased by the same amount to account for enhanced rates in the presence of minerals. According to [62] the rate of addition of an incoming G monomer across a C nucleotide on the template is 5.6 *h*^−1^ in presence of mineral Iron(II), which is 3.54 times the rate measured in [35] in the absence of minerals. Hence we multiply all of the rates taken from [35, 41] with a factor of 3.54 to get Table-1 and Table-2. The strands which do not act as templates because of their lower free energy, fold into secondary structures in the lower temperature of the semi-wet phase.

**Table 1:** Primer extension rates (*h*^−1^) in the presence of minerals, after a match in the previous step

| Templating nucleotide | Incoming monomer |  |  |  |
| --- | --- | --- | --- | --- |
|  | A | U | G | C |
| A | 0.248 | 1.878 | 0.603 | 0.284 |
| U | 3.261 | 0.425 | 1.347 | 0.06 |
| G | 0.142 | 1.701 | 0.567 | 32.005 |
| C | 0.06 | 0.074 | 5.6 | 0.078 |

**Table 2:**
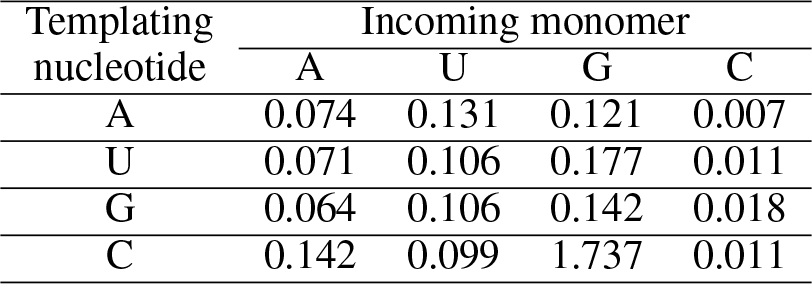
Primer extension rates (*h*^−1^) in the presence of minerals, after a mismatch in previous step

In the wet phase, the templates and their respective primers remain connected by hydrogen bonds. The templates whose primers are not fully extended have a dangling part prone to hydrolysis. The rate of such process is proportional to the number of phosphodiester bonds 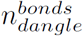 in the dangling portion i.e. the rate will be 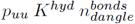, where the prefactor *p*_*uu*_ = 1.0 for the dangle. The bonds of the double stranded regions of template-primer pairs are less prone to hydrolysis as those bonds have no open *OH*^−^ groups that water molecules can attack. Hence, for the paired regions, the hydrolysis rate is given by 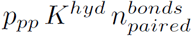, where we take prefactor *p*_*pp*_ = 0.01 for the paired region. The phosphodiester bonds of folded single strands are also susceptible to hydrolysis. But the folded (secondary) structures of such sequences, that promote base pairing between different regions of the sequence, can protect them from hydrolysis in the same way as the paired region of a template-primer is protected. Here also, the hydrolysis rate of a bond depends on whether the bases on both side of the bond are paired or unpaired. If both of them are paired then the prefactor is *p*_*pp*_ = 0.01. If both of them are unpaired, the prefactor is *p*_*uu*_ = 1.0. And if one of them is unpaired, then the prefactor is taken to be *p*_*up*_ = 0.1. Figure-2 shows different types of phosphodiester bonds, depending on the paired or unpaired nature of their neighboring nucleotides. We take the maximum hydrolysis rate per bond *K*^*hyd*^ = 0.04 *h*^−1^ from experiments [66]. Both folded strands and template-primer pairs can get hydrolyzed into multiple short fragments in the wet phase. But if a hydrolyzable bond is located in the protected regions, the breakage of that bond does not immediately result in the formation of two separate strands. They are separated from the paired region only at the time of transition between the wet and dry phase, when the temperature rises up and breaks down all hydrogen bonds of the template-primer pairs and the folded strands. Thus, at the beginning of the dry phase, all templates get separated from their respective primers, the folded single strands also unfold and the system then consists of many single strands of shorter lengths. In the wet phase the polymers become more diffusive compared to the dry and semi-wet phase. But their diffusion coefficient depends on the number of nucleotides in the polymer according to the relation *D* ~ *L*^−*γ*^, where *γ* = 0.588 [67]. Following this relation we assume that the template-primer pairs and the folded strands still have low diffusion coefficient in the wet phase, as they have many nucleotides and do not immediately fragment due to the protection provided by the hydrogen bonds. Their number density also remain low, which further reduces their diffusivity. But after they are broken up at the transition time, the shorter strands become more diffusive than their precursors (assuming water does not dry up immediately at the transition time) and their number density also increases. Hence, those shorter strands diffuse out of the system volume if there is a high concentration of short strands at the transition time. Effectively, this amounts to modeling new RNA strand formation in a localized region in space. In order to avoid instabilities associated with exponential growth in the number strands, we use a typical of re-sampling technique by defining *N*_*max*_ as the maximum number of strands that we track in the code. This number is chosen to be large enough to be representative of the population. After hydrolysis, if the number of strands becomes more than *N*_*max*_, we sample out *N*_*max*_ strands from them with probabilities (1 – *L*^−*γ*^) following the relation *D* ~ *L*^−*γ*^. The remaining strands are assumed to have diffused out of the system volume.

During the dry phase, all single strands can undergo spontaneous concatenation by step-wise addition of free monomers at their 3’ ends. The concatenation rates for addition of each type of monomer are taken to be equal as the reaction yields using different types of monomers are found to be of nearly the same order [64] and given by *K*^*con*^ = 0.62 *h*^−1^ (which is the concatenation rate of two D-type activated Adenosine nucleotides). New dimers can also form in the dry phase with the same rate. Spontaneous ligation between polymers is neglected as stated earlier. After the dry phase as the temperature drops, the next wet or semi-wet phase (depending on the model under discussion) begins and all of relevant processes are repeated after each 24-hour cycle. We use standard Gillespie algorithm [71] for carrying out the various types of reactions. The secondary structures and free energies of the RNA strands are derived using the ViennaRNA package [70].

### 2.1 Classification and detection of secondary structures

The secondary structure in Dot-Bracket notation is derived from the ViennaRNA package. The dots indicate unpaired bases. The round brackets: (and) together indicate that two bases are paired to each other. To classify the different secondary structures that emerge, we ignore the unpaired bases and remove them from the Dot-Bracket structures. We developed an algorithm for detecting 4 types of secondary structures based on the arrangement of the open and closed brackets. The types of secondary structures considered are: single-hairpin, double-hairpin, hammerhead and cloverleaf. Figure-3 shows these 4 types of secondary structures.

- **Single Hairpin:** A hairpin loop will be indicated by a set of consecutive open brackets: (, followed by a set of consecutive close brackets:) in equal numbers. We find the sets which contain repeated brackets, either open or close. If the number of such sets is 2, where the first one contains open brackets and second one contains close brackets and the number of brackets in both sets is equal then a Single Hairpin loop is detected.
- **Double Hairpin:** If the number of sets is 4, where the first and the third set contain open brackets and the second and fourth set contain close brackets, and the number of brackets in the first & second set and in the third & fourth set are equal, then a Double Hairpin loop is detected.
- **Hammerhead:** If the number of sets is 4, where the first & third set contain open brackets and the second & fourth set contain close brackets, but the number of brackets in first & fourth set is more than the number of brackets in second & third set respectively, and the condition #(first - second) = #(fourth - third) is satisfied, then a Hammerhead structure is detected.
- **Cloverleaf:** If the number of sets is 6, where the first, third and fifth sets contain open brackets and the second, fourth and sixth sets contain close brackets, with the number of brackets in the first & second set, in the third & fourth set and in fifth & sixth set being equal, then a Cloverleaf structure is detected.

**Figure 3:**
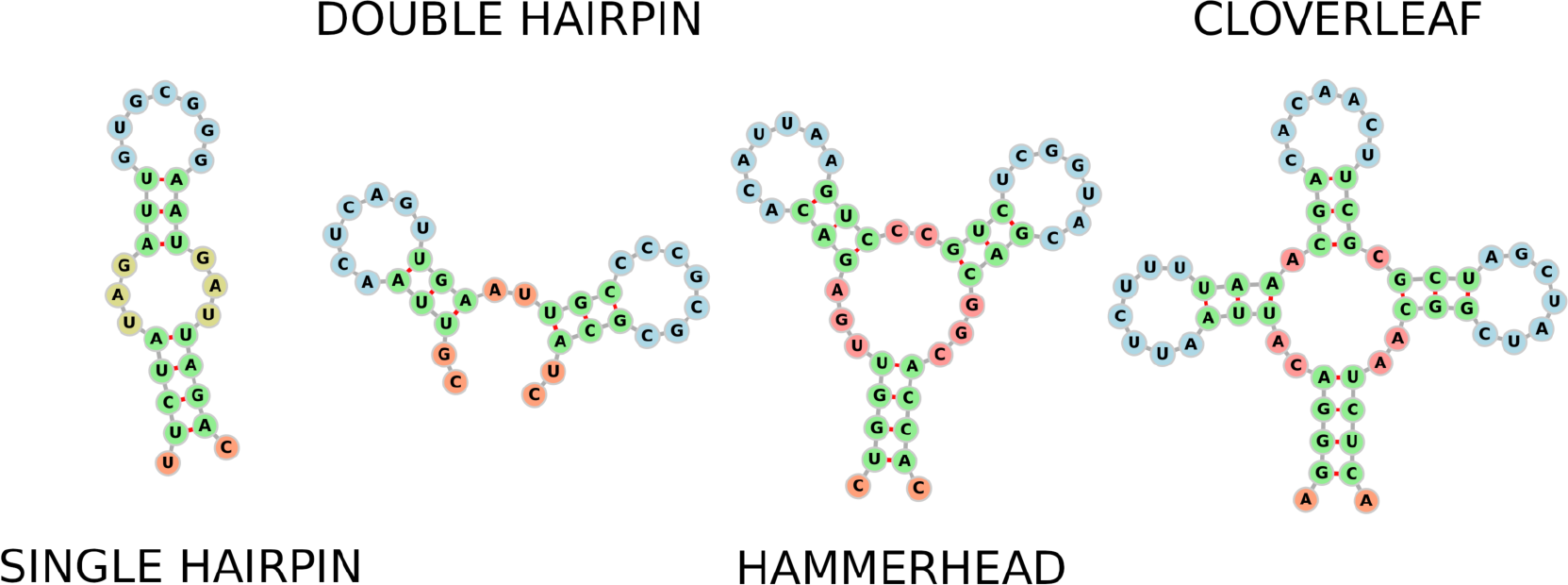
Classification of secondary structures. Blue indicates hairpin loops. Green indicates stems. Yellow indicates internal loops. Pink and orange indicate other unpaired nucleotides.

## 3 Results

### 3.1 Two-phase model: Analytical results and numerical simulations

In this section we develop a simplified analytical model for a system with 2 phases (dry and wet). In the dry phase dimerization and concatenation reaction occur just as in the main model; but instead of four types of monomers, we have only one type of monomer, with concatenation rate that is four times the rate for each type of monomer taken
previously i.e. *K*^*con*^ = 2.48 *h*^−1^. In the wet phase polymers can break up due to hydrolysis, but the hydrolysis rate for each phosphodiester bond is taken to be equal in this case. We assume that in the wet phase polymers can totally degrade into its constituent monomers at a fixed rate *K*^*d*^. As in the main model, the reaction rates are considered to be independent of monomer concentration. We use a mean field approach to solve this problem analytically. Two parameters which can change because of the reactions, are the total number of polymers (*N*_*T*_) and the total length of all polymers (*L*_*T*_).

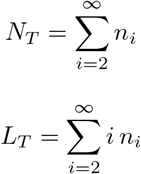

Here *n*_*i*_ is the number of polymers containing *i* nucleotides. First we consider a system with a single phase, where all reactions occur simultaneously. As a result of the concatenation reaction between two monomers (i.e. dimerization), a new dimer can get created at a rate *K*^*dim*^. Although for constant monomer concentrations, dimerization is a zeroth order reaction and concatenation of strands of length ≥ 2 are first order reactions; it is reasonable to associate very similar rates to these processes in our mean field model, because the rates of these processes are found to be similar in experiments as well [64, 72]. We write *K*^*dim*^ = *αK*^*con*^, where *α* is a correction prefactor that determines the extent to which the concatenation and dimerization rates differ. Concatenation between a polymer and a monomer does not create a new polymer, it only increases the length of the polymer by one nucleotide. Hence, *N*_*T*_ will increase because of dimerization at a rate *α K*^*con*^ and *L*_*T*_ will increase due to both concatenation and dimerization reactions at a rate *K*^*con*^(*N*_*T*_ + 2*α*) respectively.

*N*_*T*_ strands contain (*L*_*T*_ – *N*_*T*_) bonds, each of which can get hydrolyzed. Out of these (*L*_*T*_ – *N*_*T*_) bonds, breaking of (*L*_*T*_ – 3*N*_*T*_) bonds will result in creation of new polymers, as breaking of the bonds at the two ends of a polymer leads to the creation of a polymer and a free monomer (which is not a polymer by definition). The total length of all polymers *L_T_* can decrease due to hydrolysis only when any of the bonds at the two ends of a polymer gets hydrolyzed.
Hence by hydrolysis *N*_*T*_ can increase and *L*_*T*_ can decrease at rates *K*^*hyd*^(*L*_*T*_ – 3*N*_*T*_) and 2*K*^*hyd*^*N*_*T*_. Finally, total degradation of polymers into its constituent monomers causes both *N*_*T*_ and *L*_*T*_ to decrease at rates *K*^*d*^*N*_*T*_ and *K*^*d*^*L*_*T*_ respectively. Hence, the mean-field equations governing the dynamics of the system are,

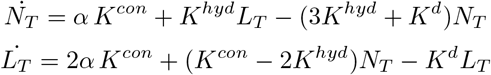

The average length 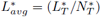 at equilibrium is,

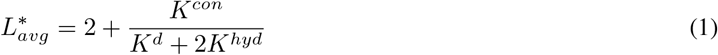

The expression shows that the contribution to the average length comes from dimers as well as the relative importance of the antagonistic processes of concatenation and degradation & hydrolysis that acts to respectively increase and decrease the size of polymers. Moreover, the prefactor *α* for dimerization rate drops out from this expression for average length because it controls only the number of polymers created and not their average size. Hence, we can safely assume that dimerization and concatenation of a monomer and a polymer happens effectively at the same rate *K*^*con*^, for our following 2-phase analytical model and 3-phase numerical simulations.

Substituting *K*^*con*^ = 2.48 *h*^−1^ and *K*^*hyd*^ = 0.04 *h*^−1^ and choosing *K*^*d*^ = 0.3 *h*^−1^ we get, 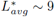 nucleotides. The equations for 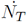 and 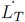 can be integrated numerically to obtain the time variation of the average length. Figure-4(A) shows the comparison between the analytically obtained average length vs time plot which clearly matches with the simulation results.

**Figure 4:**
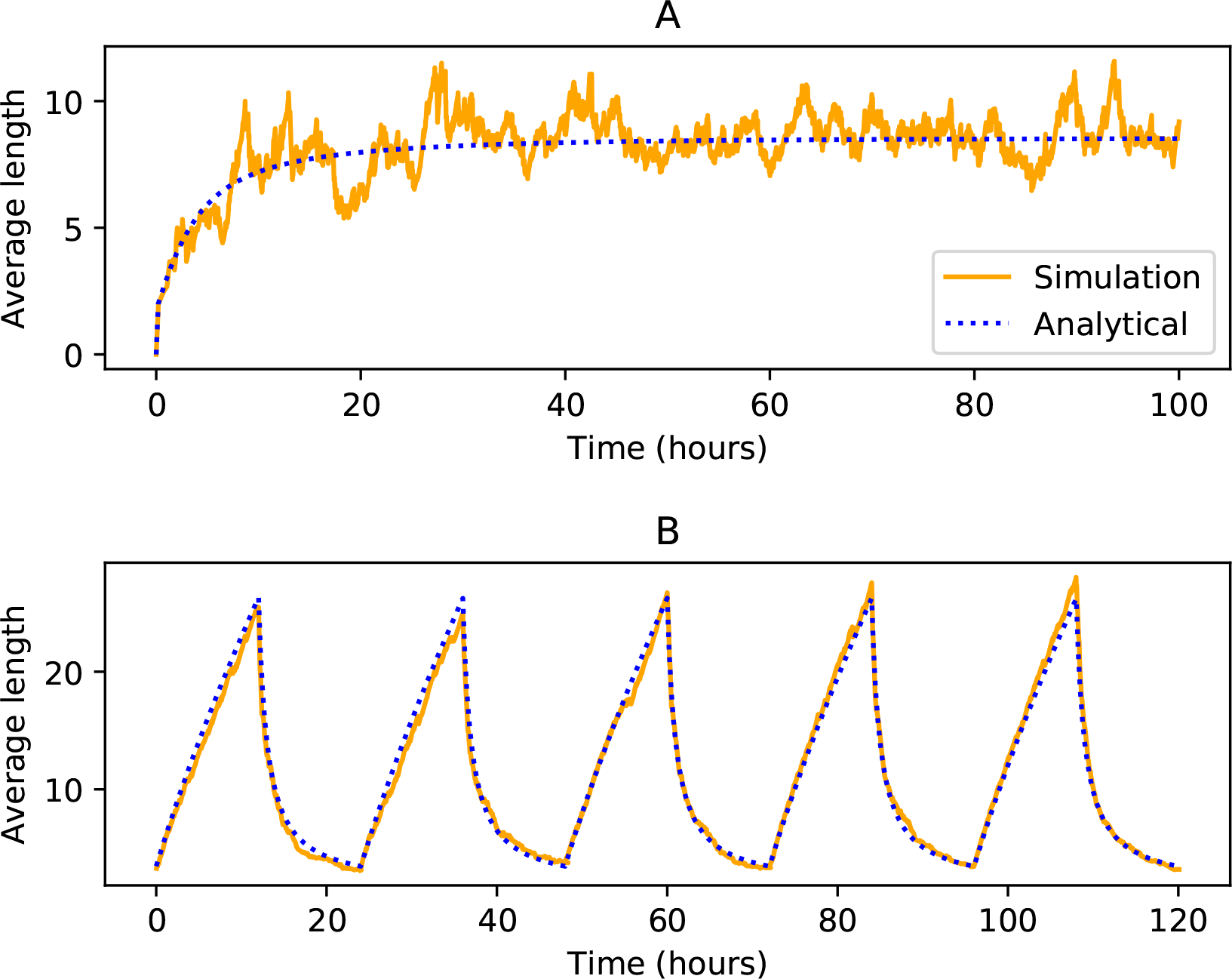
Time evolution of the average polymer length (in number of nucleotides) from both analytical calculations (blue dotted line) and numerical simulation (orange solid line) for the case when **A:** without cycling when all reactions occur simultaneously; **B:** with cycling when each day is divided into a dry and wet phase of 12 hours each. The plot shows the variation over a period of several days. Other parameters used: *K*^*con*^ = 2.48 *h*^−1^; *K*^*hyd*^ = 0.04 *h*^−1^; *K*^*d*^ = 0.3 *h*^−1^ in (A) and *K*^*d*^ = 0.2 *h*^−1^ in (B).

We then model environmental cycling effects by separating the dry and wet phases in the system. The equations for 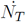 and 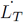 will be now different for the different phases. In the dry phase only the concatenation reactions occur whereas the hydrolysis and degradation reactions take place only in the wet phase. Hence, the following equations will now govern the system dynamics.

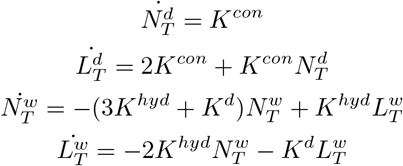

Here the superscript **d** and **w** denote the dry and wet phase respectively. In the dry phase the coupled equations for 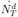 and 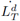 can be easily solved to give,

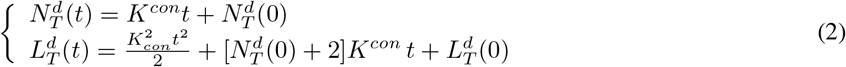

In the wet phase the coupled equations can be written in matrix form as,

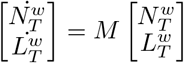

with

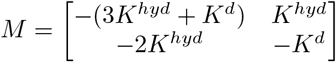

The matrix M has eigenvalues *λ*_1_ = −2*K*^*hyd*^ – *K*^*d*^ and *λ*_2_ = −*K*^*hyd*^ –*K*^*d*^ with corresponding eigenvectors (1,1) and (1/2, 1). Hence, in the wet phase, the total number of polymers and total length of all polymers vary as,

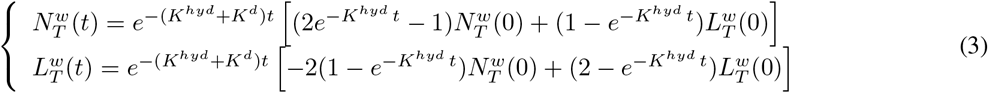

In equilibrium, the cyclic boundary conditions are:

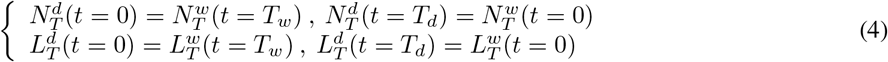

where *T*_*d*_ and *T*_*w*_ are the duration of the dry and wet phase. Applying these cyclic boundary conditions and using the dry phase solutions (Eq. 2) for 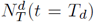 and 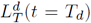 leads to a pair of linear equations in 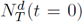 and 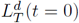. Solving them numerically allows us to obtain exact analytical expressions for the time evolution of the total number and length of polymers.

For example, using *K*^*con*^ = 2.48 *h*^−1^, *K*^*hyd*^ = 0.04 h^−1^, *K*^*d*^ = 0.2 h^−1^ and *T*_*d*_ = *T*_*w*_ = 12 h we get, 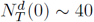 and 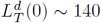. After determining the initial values of *N*_*T*_ and *L*_*T*_ at the beginning of each dry and wet phase, it is easy to get the equilibrium time variation of the average length *L*_*avg*_(*t*) = *L*_*T*_ (*t*)*/N*_*T*_ (*t*) for the dry and wet phase from Eq. 2 and Eq. 3 respectively.

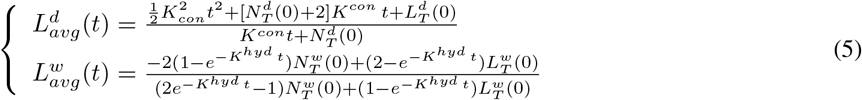

Figure-4(B) shows the time variation of the average length over 5 days at equilibrium, obtained from the analytical model which match perfectly with the results of simulations. It is also clear from comparing the two models that the maximum length of strands in the 2-phase model is larger than the maximum length obtained in the single phase model as long as the duration of the dry phase is above a certain threshold. For the set of parameters used to generate Figure-4(A), this threshold dry phase duration is ~ 5 hours. Our 2-phase model reinforces the importance of environmental cycling in increasing the length of polymers generated.

Using the analytical model we can also study the effect of non-templated ligation of RNA strands. Upon inclusion of spontaneous ligation in the analytical model we find that (see *Supplementary information*), ligation affects the average length of strands only when *K*_*lig*_ ~ *K*_*con*_. Since we know from experiments that *K*_*lig*_ = 10^−3^ *K*_*con*_ [64, 68], we can justify neglecting spontaneous ligation of RNA strands in the three-phase model described in the next section.

### 3.2 Three-phase Model: Effect of the duration of different phases

In this section, we investigate the role of non-templated concatenation as well as template-directed primer extension processes in the formation of long sequences with complex structures as the duration of the three environmental phases are varied. Formation of long RNA polymers is a key prerequisite for the emergence of complex structures. Quantities like the average length, the maximum length and average free energy of synthesized strands are useful metrics that determine the fraction of templates in the population and relative abundance of the four types of secondary structures depicted in Figure-3. The three environmental phases (dry, semi-wet and wet) are distinguished by the reactions they support and hence we varied the duration of these 3 phases to gauge the impact of different chemical processes on the emergence of complex secondary structures. Since the length of a day was fixed at 24 hours, we chose to independently vary two (dry and semi-wet) of the three phases which automatically constrains the duration of the remaining (wet) phase.

Figure-5 shows the time evolution of the average length of strands, average free energy (which can be used as a proxy for folding efficiency of the strands) at the end of the dry phase and the average length of primers at the end of the semi-wet phase, for different duration of the three phases. We also obtained 2D heat maps (Figure-6) for the time averaged values (obtained by time-averaging over 100 days after equilibration) of the maximum length (Figure-6(A)) and average free energy (Figure-6(B)) of the strands and maximum length of templates (Figure-6(C)) and primers (Figure-6(D)) that allowed us to compare in greater detail the impact of each phase duration on these quantities. The length distribution of strands for different duration of the three phases are shown in Supplementary Figure-S2. We observe an increase in average length [Figure-5(A) and Supplementary Figure-S1(A)] and maximum length (Figure-6(A)) of strands and decrease in average free energy (Figure-5(B) and Figure-6(B)) with increase in duration of both dry and semi-wet phase. Increase in the duration of the dry phase allows the concatenation process to increases the length of the strands and their folding efficiency which in turn protects them from hydrolysis due to their secondary structures. Hence, we get longer strands with lower free energy, with an increase in the duration of the dry phase. For a fixed duration of the dry phase, an increase in the duration of the semi-wet phase also leads to increase in the maximum and average length of the strands as can be seen from the columns of Figure-6(A) and Supplementary Figure-S1(A).This is because the increased duration of the semi-wet phase implies a shorter duration of the wet phase that suppresses the break-up of long strands by reducing the likelihood of hydrolysis. Moreover, an increase in the duration of the semi-wet phase also gives the templates more time to create longer primers with more structural complexity due to increased mis-incorporation during primer extension. Longer primers imply templates with shorter dangles, which will reduce their hydrolysis rates as greater fraction of the template-primer pair will be paired and hence protected. Figure-5(C) shows the time evolution of the average length of primers for different duration of dry and semi-wet phase. The maximum length of templates increases in a similar fashion with increase in the duration of dry and semi-wet phase [Figure-6(C)]. Higher average and maximum length of strands for longer duration of the dry phase and a non-zero fraction of templates [Supplementary Figure-S1(B)] in the system implies that the chosen templates will be of higher lengths as well. Hence, the maximum length of templates also increase with increasing duration of the dry phase. Increase in the semi-wet phase duration in this case helps indirectly by reducing the wet phase duration, which in turn boosts the average and maximum length of strands, thereby increasing the maximum length of templates as well. We have also verified that as the dry phase duration increases relative to that of the other phases, the concatenation process starts dominating and the effectiveness of the template-directed primer extension process in increasing the average length gradually decreases to zero.

**Figure 5:**
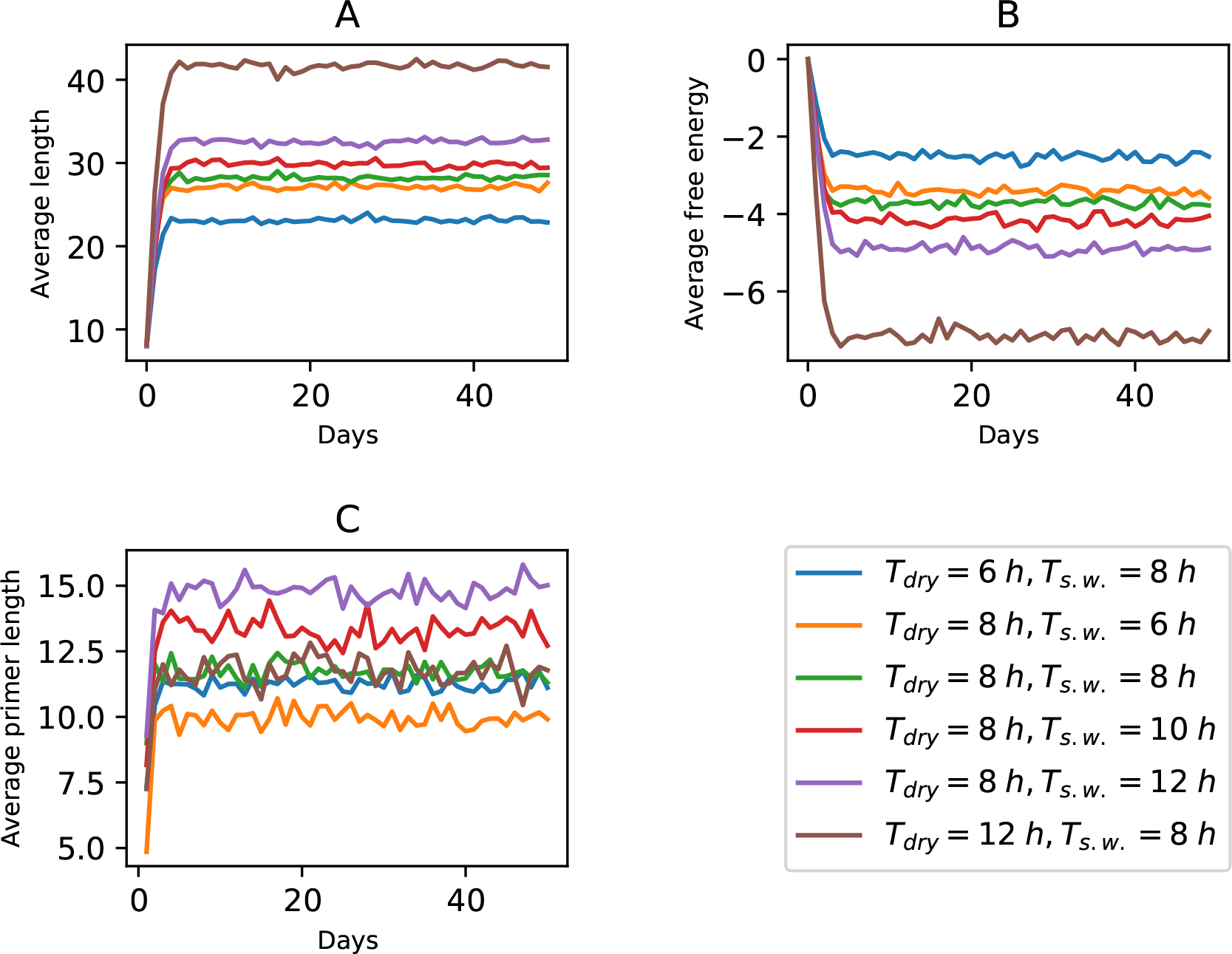
Time evolution of the **A:** average length of the strands, **B:** average free energy of the strands, measured in kcal/mol and **C:** average length of primers, for six different duration of the dry and semi-wet phases.

**Figure 6:**
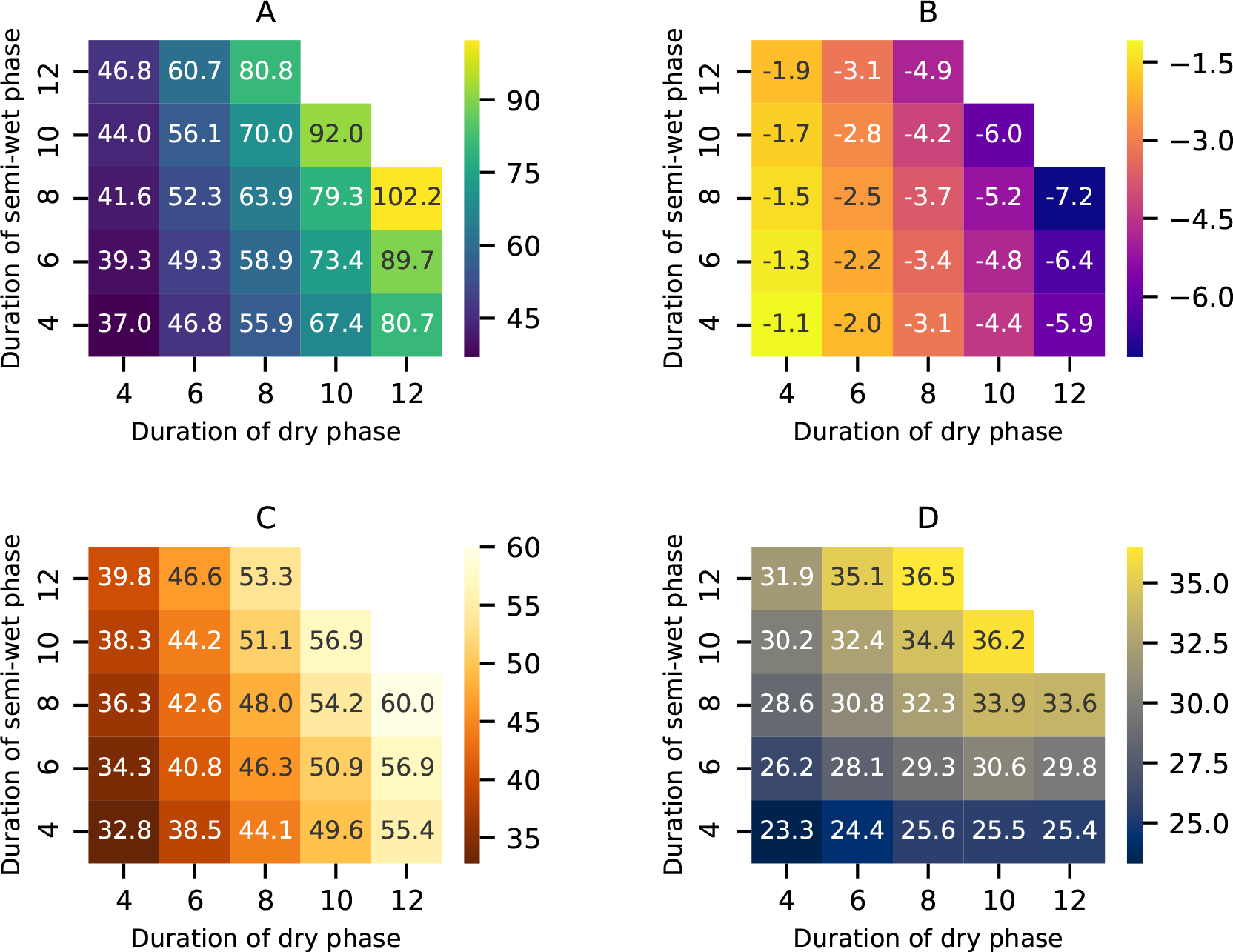
**A:** Maximum length of RNA strands and **B:** average free energy for folding of RNA strands at the end of the dry phase, **C:** maximum template length and **D:** maximum length of primers; for different duration of dry and semi-wet phase, obtained by averaging over 100 days after equilibration.

It is clear from Figure-6(A) and supplementary Figure-S1(A) that a lower average free energy (compare with Figure-6(B)) is clearly correlated with larger average and maximum length of the strands which in turn increases with the duration of the dry phase. However, the average and maximum length of the primers does not show any significant increase with increase in duration of the dry phase for fixed duration of the semi-wet phase [see Figure-6(D) & Supplementary Figure-S1(C)] but increase more rapidly with increase in the duration of semi-wet phase. In most of the cases, the average length of primers is greater than 10 nucleotides with the maximum length in several cases extending well beyond 30 nucleotides under favourable environmental conditions. This signifies that the template-directed primer extension is effective in creating long complementary replicates despite the slowdown of the primer extension rates after mismatches.

The fraction of templates (i.e. those strands possessing the ability to create complimentary copies) decreases with increase in the duration of both dry and semi-wet phase [Supplementary Figure-S1(B)] because increasing the duration of these phases favors longer strands with better folding efficiency. Hence, more strands fold leaving fewer strands to act as templates. Nevertheless, the minimum fraction obtained is 0.15, which is still quite significant to facilitate template-directed primer extension processes. The fraction of templates also increases with increase in average free energy Figure-6(B) as expected, since sequences with larger average free energies are less likely to fold into complex secondary structures.

In Figure-7 we checked how the abundance of different secondary structures vary with the duration of dry and semi-wet phase. The abundance of single hairpin structures (Figure-7(A)) increase with increase in the duration of dry and semi-wet phase up to moderate values, but for even longer duration of those phases we see a decrease in the abundance of single hairpins. The abundance of the double hairpin, hammerhead and cloverleaf structures [Figure-7(B), (C), (D)] on the other hand always increase with increase in the duration of dry and semi-wet phase. For lower duration of the dry phase, the length of sequences produced are not large enough to permit the formation of such complex secondary structures which require polymer lengths larger than ~ 45 (for hammerhead structures) or ~ 50 (for cloverleaf structures). Increasing the duration of the semi-wet phase for fixed duration of the dry phase can also facilitate the emergence of more complex secondary structures as observed from [Figure-7(C),(D)]. This occurs due to the lower likelihood of breaking up of larger existing sequences via hydrolysis as well as the increased time available for replication via template-directed primer extension. The former factor also explains the increase in maximum length of sequences observed along a column in (Figure-6(A)) while the latter factor can also lead to increased structural complexity through incorporation of mismatches during replication via the primer-extension process. An increase in duration of the semi-wet phase coupled with a decrease in duration of the wet phase leads to the increase in the number of longer polymer fragments at the beginning of the dry phase each of which can be further extended through the non-templated concatenation process prevalent in the dry phase. In summary, a short to medium duration of dry and semi-wet phase favor single hairpin structures, but longer dry and semi-wet phases promote emergence of more complex structures thereby underscoring the importance of both non-templated as well as template-directed polymer extension processes.

**Figure 7:**
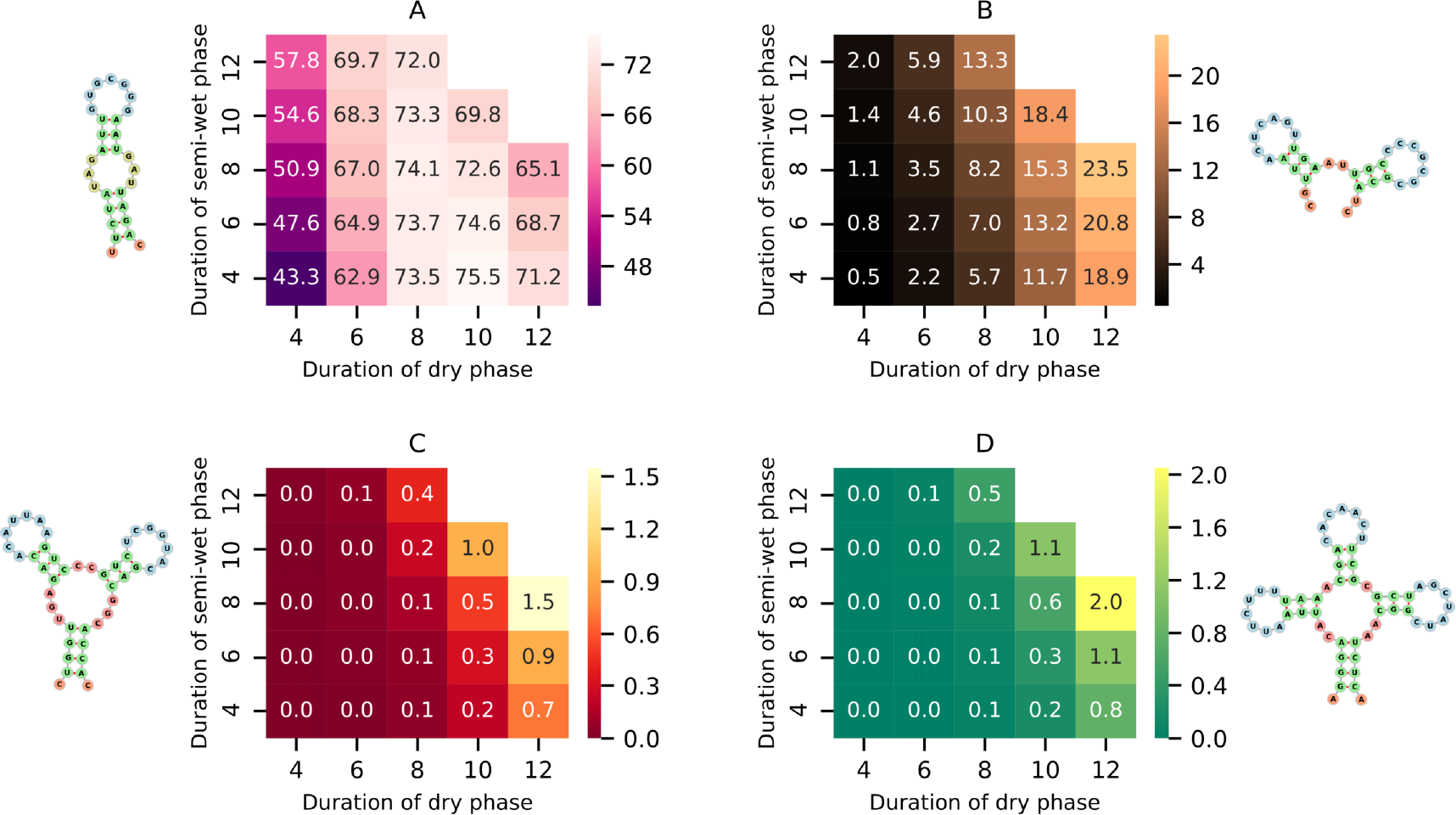
Average (percentage) abundance of **A:** single hairpin structures, **B:** double hairpin structures, **C:** hammerhead structures and **D:** cloverleaf structures; at the end of the dry phase for different duration of dry and semi-wet phase.

The average percentage of mismatch in the primers of length greater than 10 nucleotides increase very slightly for increase in the duration of the dry phase. This can be attributed to the production of longer templates increasing the likelihood of mismatches even though the semi-wet phase duration remains fixed. But the percentage error increases rapidly with increase in the duration of the semi-wet phase [Supplementary Figure-S1(D)]. From Table-2 it is evident that the primer extension rates decrease significantly after a mismatch. Hence the primers can extend to greater lengths only if they have fewer number of mismatches. For small duration of the semi-wet phase only those primers which have fewer number of mismatches, will extend their lengths by a large amount, as the primers get a relatively small amount of time to extend. But for higher duration of the semi-wet phase, even primers with more number of mismatches can also significantly increase their lengths as all primers then get more time to extend. Hence, a smaller duration of the semi-wet phase favors error free replication of strands.

The hydrolysis and non-templated concatenation rates are not known as accurately as template-directed extension rates. To account for the possibility of variation of these rates, we carried out simulations where these rates as well as the protection prefactors of the phosphodiester bonds were varied over a plausible range. Figure-8(A) shows that increasing the prefactors reduces the average length of the strands, up to the theoretical limit when all bonds are equally susceptible to hydrolysis. Reduction in the hydrolysis rate causes increase in average length (Figure-8(B)). But increase in hydrolysis rate even by a couple of orders of magnitude does not cause significant decrease in the average length because we calculate the average length at the end of dry phase. Hence, even if the high rate of hydrolysis causes total breakdown of strands into monomers, new strands are created and extended in the dry phase by dimerization and concatenation processes. This leads to a lower bound on the length of RNA strands that depends only on the concatenation rate. Figure-8(C) also shows that the average length is highly dependent on the concatenation rate.

**Figure 8:**
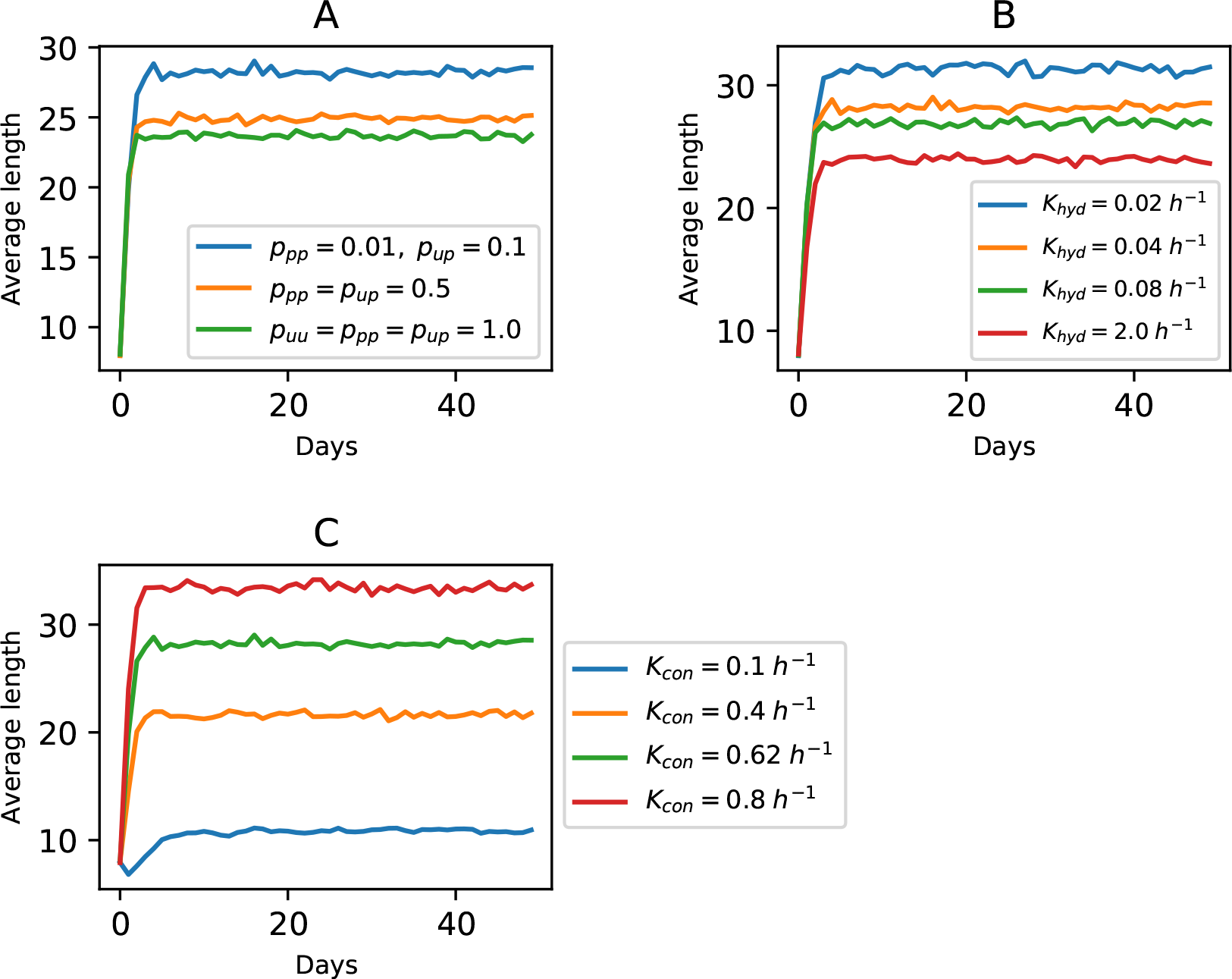
Average length of strands vs time for **A:** different values of protection prefactors, **B:** different hydrolysis rates and **C:** different concatenation rates; when each of the 3 phases are of duration 8 hours

## 4 Conclusion

The ability to generate long sequences with complex structures that encode catalytic functions, through non-enzymatic processes acting on basic chemical building blocks, is the holy grail of the RNA world scenario. Our results show that by using realistic reaction rates derived from experiments, it is possible to generate long RNA strands in plausible primordial environmental conditions that allow for temporal segregation of reactions that promote non-templated concatenation, template-directed primer extension and hydrolysis. The efficacy of evolving structurally complex sequences therefore crucially depend on the duration of the dry and semi-wet phases. Despite the relatively low rate of the non-templated concatenation reactions, they are crucial for extending the length of the polymers in the existing pool thereby increasing the likelihood of generating longer templates. This in turn facilitates replication through the faster template-directed primer extension process.

An alternative model to the one examined here involves self-assembly of a complete ribozyme from constituent sequence fragments [73, 74]. Recent work has shown that larger ribozymes (≳ 200 nucleotides) can be synthesized from smaller constituent fragments each of which catalyze the formation of another fragment in a cyclic manner. Such auto-catalytic hypercycles [75] have a selective advantage [74] over selfish replicators that catalyze their own assembly from substrate molecules [74, 76, 77]. However, such a model still needs to explain how large sequence fragments first appeared. The prebiotic processes discussed here provide a mechanism for synthesizing the sufficiently large sequence fragments that can form the nodes of a hypercyclic network needed to cooperatively assemble larger ribozymes like the *Azoarcus* ribozyme.

In our model we assumed that all template-primer pairs get separated from each other in the dry phase, which has a maximum temperature of ~ 90° *C*. But in reality, the template-primer pairs with a more than ~ 30 paired nucleotides (even with maximum ~ 30% mismatched nucleotides) will not get separated in the dry phase, as their melting temperatures are higher than 90° *C* [78, 79]. For such pairs even if some of the phosphodiester bonds in the paired regions get hydrolyzed in the wet phase, those bonds will re-form as these template-primer pairs will still remain connected in the dry phase. Hence, these template-primer sets will remain paired in the following day, undergo further extension of the primers in the next semi-wet phase and can eventually get fully extended. This will lead to an increase in the probability of formation of complementary template-primer pairs with more than 30 paired nucleotides, which are more resistant to strand separation in the hot, dry phase. In this way stable double stranded RNA molecules can form. From Figure-6 we see that the maximum length of primers is greater than 30 nucleotides for several different duration of the dry and semi-wet phase. The more stable double stranded RNA molecules that emerge will serve as reliable storage of genetic information, whereas some of the ribozyme-like structures resulting from folded single strands may start showing enzymatic activity. Once that happens, some of the functional RNA molecules (e.g. replicases) may catalyze the process of error free replication by reducing the rates of mis-incorporations or mutations. However, the effectiveness of the increased duration of the dry phase in generating longer sequences is incumbent on the unlimited supply of monomers for non-templated concatenation process. While this is a reasonable assumption as long as the duration of the dry phase is finite, this becomes less and less valid as the duration of the dry phase becomes very long. Hence, periodic hydration is required to allow for replenishment of the monomer pool and keep the concatenation reaction independent of monomer concentration. Moreover, we see from Figure-7(C) & (D) that increasing the duration of the semi-wet phase for moderate duration of the dry phase can also help in generating long sequences with complex secondary structures. Hence, the presence of the 3 phases creates the most ideal environment for functional bio-molecules to emerge in an RNA world.

The transition between any two phases is very sharp in our model. But in reality the transition between the phases is likely to be continuous. During the transition between dry and semi-wet phase, while the temperature drops gradually, the sequences with lower free energies will fold before the ones with comparatively higher free energy. Such an overlap between the dry and semi-wet phases also amount to coexistence of the concatenation and template-directed primer extension processes. The strands which fold later will undergo further concatenation reaction, increasing their length. Primers will also start attaching to the strands that are yet to fold. Upon attachment of a primer, the primer can extend along the 3’ – 5’ direction of the template by template-directed primer extension while the template can also extend along the 5’ – 3’ direction by concatenation. Hence coexistence of concatenation and template-directed primer extension processes can lead to template-primer pairs having dangles on both sides, with the central portion being connected by hydrogen bonds. Continuous transition between semi-wet and wet phase will lead to coexistence of template-directed primer extension and hydrolysis which act antagonistically. On one hand, the primer extension process can then take place for some duration giving more time for the primers to extend fully. However, that will be possible only when the templates do not get truncated by the onset of hydrolysis towards the end of the semi-wet phase. The average length of sequences would then be modulated by the relative importance of these competing processes. At the transition between wet and dry phase the gradual increase of temperature can result in the appearance of a semi-wet phase at their transition time, which will give further boost to the full extension of those primers that don’t get separated from their templates due to long paired region. Hence continuous transitions will likely favor the emergence of double stranded regions in the RNA molecules. This, in turn, could protect them from back hydrolyzing completely, thus leading to a further increase in the maximum length of the resulting RNA strands in these realistic scenarios.

Our model assumes a continuous and constant supply of all type of monomers during all three phases, implying that each type of monomer has a fixed and equal concentration in the system. But it is quite possible that there is greater supply of monomers in the wet phase due to high diffusivity and consequently lower supply of monomers in the dry and semi-wet phase i.e. besides preventing polymer diffusion, dry conditions are likely to inhibit monomer diffusion too. Then the monomer concentration will not remain constant with time and the spontaneous and template-directed primer extension rates will depend on the monomer concentration at any instant. But even in that case, the emergence of complex structured RNA and double stranded RNA is possible if the total number of monomers in the system is significantly higher than the total number of strands; which implies that the change of monomer concentrations will be negligible, thereby ensuring that the rates do not drop significantly.

Remarkably, our simulations generate tRNA-like secondary structures, thereby indicating that such pre-tRNA’s may have evolved quite early in the prebiotic RNA world. These precursors of tRNA’s could have eventually evolved to self-catalyze aminoacylation, non-specifically at first, but subsequently with increasing specificity brought on by selection pressures. It seems plausible that the components of a primitive translation machinery, such as pre-tRNA self-catalyzing their aminoacylation and ribozymes catalyzing amino-acylation of tRNA’s, may have progressively evolved in an RNA world [80]. Our work indicates that a longer duration of dry and semi-wet phases in an ecological niche characterized by mineral-rich muddy pools can provide the ideal conditions for emergence of long RNA strands with complex structures and potential catalytic functions. Such an RNA world will also include double-stranded RNA molecules capable of storing genetic information like DNA and therefore showcase all the characteristics that can kick-start primordial life based on RNA only.

## Acknowledgment

This work was supported in part by a CNRS PICS grant 07501 to J. Derr and S.Rajamani. S. Roy is supported by an INSPIRE graduate fellowship given by the Department of Science & Technology, Government of India.

