## Supplementary Material for "Emergence of ribozyme and tRNA-like structures from mineral-rich muddy pools on prebiotic earth"

### Supplementary information

#### 1 Supplementary figures

Figure S1: **A:** Average length of RNA strands at the end of the dry phase, **B:** fraction of templates in the system, **C:** average length of the primers **D:** average percentage of mismatches in the primers; for different duration of dry and semi-wet phase.

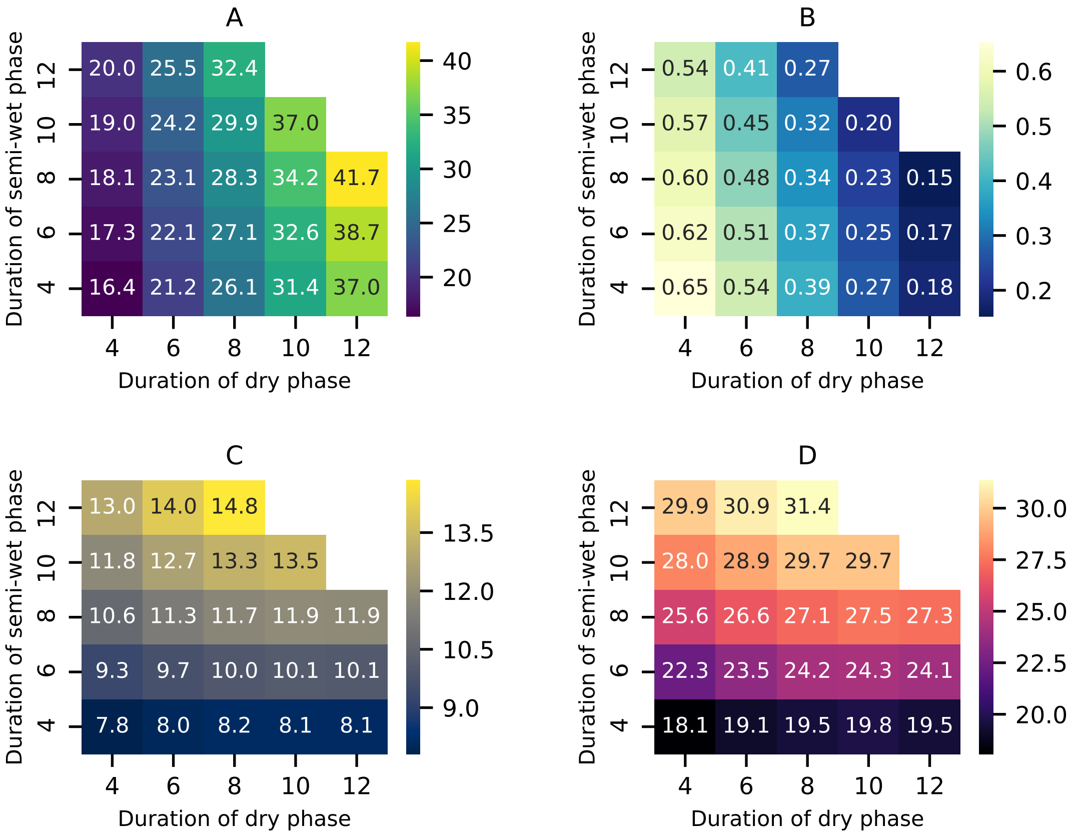

Figure S2: Length distribution of RNA strands in equilibrium when the duration of the 3 phases are **A:** dry = semi-wet = wet = 8 (hours); **B:** dry = 12, semi-wet = 8, wet = 4 (hours); **C:** dry = 8, semi-wet = 12, wet = 4 (hours) and **D:** dry = semi-wet = 6, wet = 12 (hours).

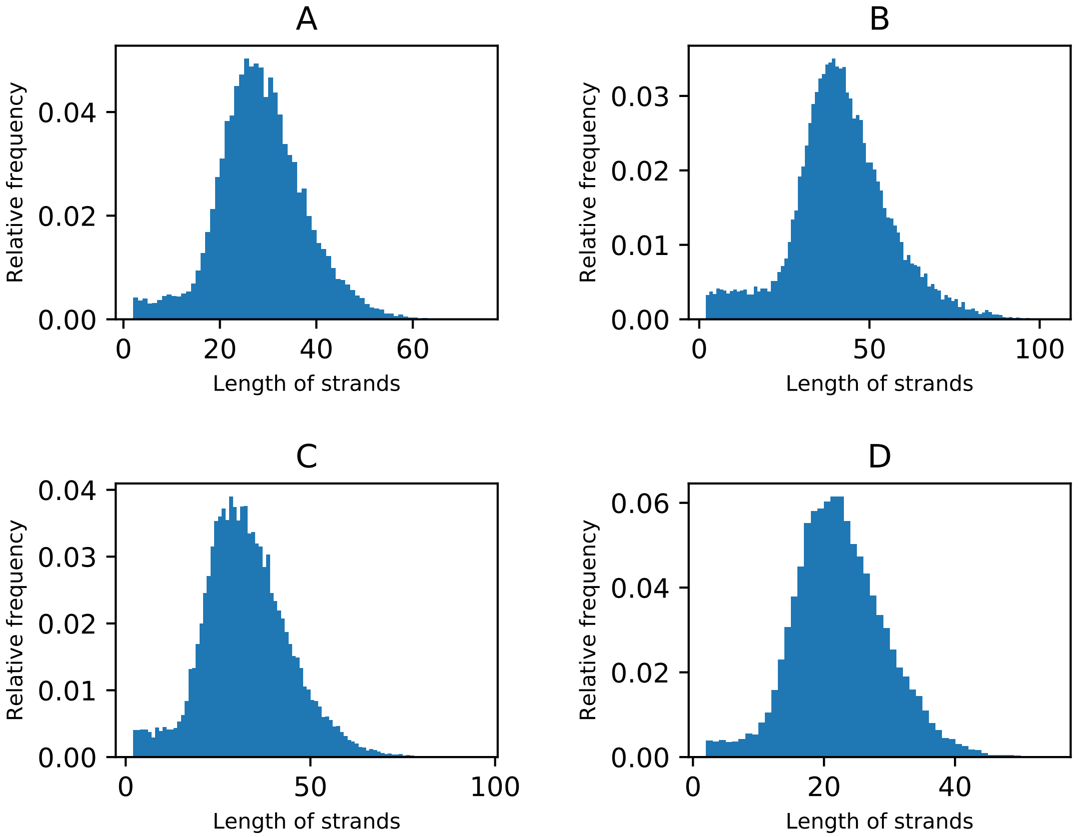

#### 2 Inclusion of spontaneous ligation processes

At lower concentration of polymers, the spontaneous ligation rate is proportional to the square of the number of strands. But for higher concentration of polymers, the ligation rate should saturate. Hence, we use a concentration depended ligation rate $K_{lig}\, tanh((N_T/N_C)^2)$. Taking this reaction into account leads to an extra term in the equation for $\dot{N_T^d}$,

$$\dot{N_T^d} = K^{con} - K_{lig}\, tanh((N_T/N_C)^2)$$

where $N_C$ is a factor that determines how quickly the function attains its extreme values. Using $N_C = 50$ we obtained the time variation of average length during any dry phase for different values of $K_{lig}$ [Supplementary Figure-S3]. It is evident from the figure that spontaneous ligation produces significant effect in the average length only when $K^{lig} \sim K^{con}$.

Figure S3: Average length vs time in dry phase in presence of non-templated ligation reaction as found from analytical model for **A:** $K_{lig} = 10^{-3}\ K_{con}$; **B:** $K_{lig} = 10^{-2}\ K_{con}$; **C:** $K_{lig} = 10^{-1}\ K_{con}$ and **D:** $K_{lig} = K_{con}$.

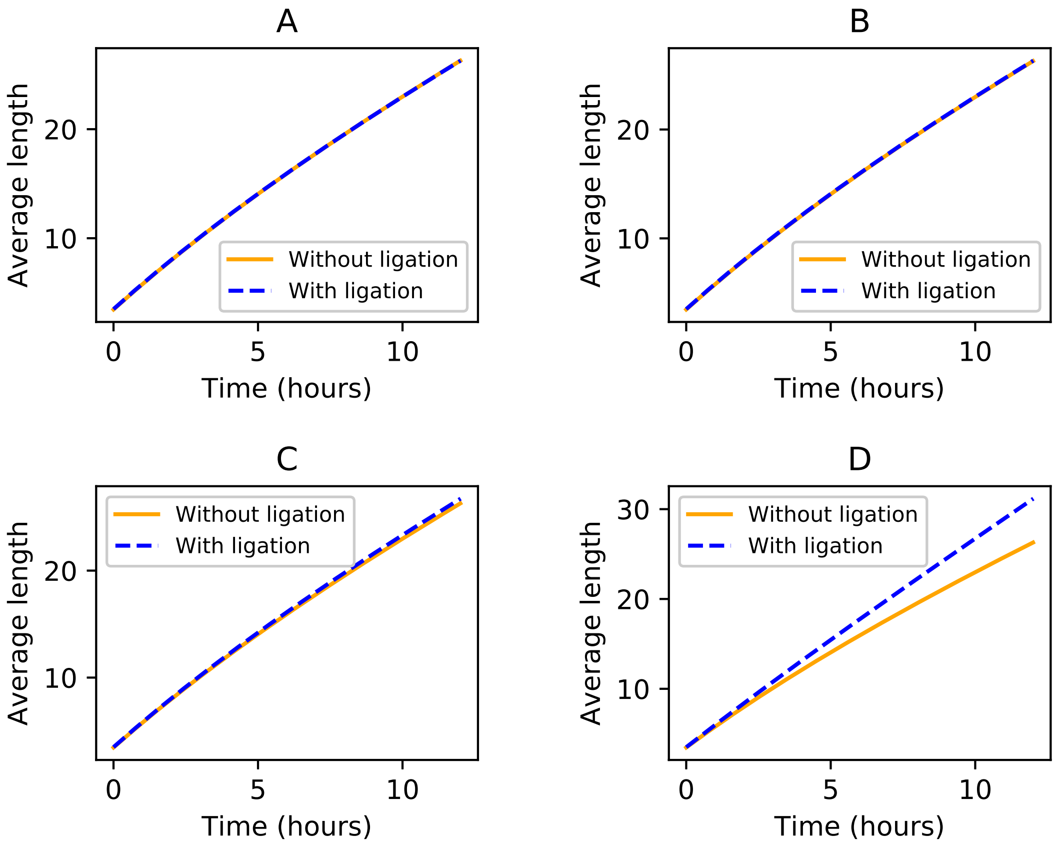

#### 3 Pseudo-code for the 3-phase simulation model

Generate $N = 100$ initial strands, of lengths normally distributed around 8 nucleotides with a standard deviation of 1 nucleotide. The initial strands are either poly-A or poly-G strands.

Begin loop over days.

**Semi-wet phase:**

1. Begin loop over set of strands
   (a) For each strand, find out it's free energy and secondary structure
   (b) Classify it's secondary structure based on the algorithm provided in the main text
   (c) Generate a random number
   (d) If the random number is $< e^{-E/E_C}$
      - Mark it as a template
2. Separate out those marked templates from the set of strands
3. The remaining strands are folded single strands
4. Begin loop over set of templates
   (a) Assign primers of length 1 nucleotide to each template
   (b) The nucleotide to be attached across the 3' end of a template is chosen randomly based on the relative reaction propensities of the 4 types of nucleotides across the 3' nucleotide (from Table-1 of main text)
5. Begin loop over the template-primer sets
   (a) For each template-primer begin loop over time up to the duration of semi-wet phase, constrained by the fact that the primer length can never exceed the length of the template
      i. Check whether the last nucleotide of the primer is complementary to the nucleotide on the template across it
      ii. If it's complementary
         - Assign primer extension rates for 4 types of monomers from the Table-1 of main text
      iii. Else
         - Assign primer extension rates from the Table-2 of main text
      iv. Draw a random-exponential time step dt based on the total rate of primer extension
      v. Add dt to the total time
      vi. If the total time after addition of dt is less than duration of the semi-wet phase
         A. Choose a nucleotide based on the relative propensities for the addition of the 4 types of nucleotide
         B. Add the chosen nucleotide to the primer
   (b) End loop over time
6. End loop over template-primer set

**Wet phase:**

1. Begin loop over template-primer sets
   (a) Begin loop over time up to the duration of wet phase
      i. Assign hydrolysis rates ($p_{uu}/p_{up}/p_{pp} * K_{hyd}$) to each phosphodiester bond of the template and primer, based on the paired or unpaired status of their neighboring nucleotides
      ii. Calculate the total hydrolysis rate $\left(= \sum_i^{bonds} p_i K_{hyd}\right)$ for this template-primer pair
      iii. Draw a random-exponential time step dt based on the total hydrolysis rate
      iv. Add dt to the total time
      v. If the total time after addition of dt is less than the duration of the wet phase
         A. Choose a bond randomly based on their relative hydrolysis rates $\left(\frac{p_i K_{hyd}}{\sum_i^{bonds} p_i K_{hyd}}\right)$
         B. If the chosen bond is in the dangle region
            - Separate the broken dangle right away and add it to population of single strands, keeping track of the time when it has broken apart from the template
         C. If the chosen bond is in the paired region of the template or primer

- Mark it as a broken bond, but don't separate right away. Separate all such bonds after the end of wet phase, when the dry phase has just begun.

(b) End loop over time

2. End loop over template-primer sets
3. Begin loop over single strands
   (a) Find out the secondary structure
   (b) Based on the secondary structure, assign hydrolysis rates ($p_{uu}/p_{up}/p_{pp} * K_{hyd}$) to each phosphodiester bond
   (c) Begin loop over time up to the full duration of the wet phase if it's a originally folded single strand
   (d) If it's a broken dangle which was added later, the time loop should start from the time when it was added to the population of single strands
      i. Calculate the total hydrolysis rate $\left(= \sum_i^{bonds} p_i K_{hyd}\right)$ of the strand
      ii. Draw a random-exponential time step dt based on the total hydrolysis rate
      iii. Add dt to the total time
      iv. If the total time after addition of dt is less the duration of wet phase
         A. Choose a bond randomly based on their relative hydrolyis rates $\left(\frac{p_i K_{hyd}}{\sum_i^{bonds} p_i K_{hyd}}\right)$
         B. Mark this bond as broken and remove from the set of intact bonds
   (e) End loop over time
   (f) Break up the strand at those marked bonds
4. End loop over single strands

**Dry phase:**

1. Separate template-primers
2. Reverse the direction of the primers (as a template and primer are directed oppositely)
3. Add the separated templates and primers to the population of single strands
4. Reject the strands which have length 1 nucleotide (as they are monomers and we don't consider monomers as strands)
5. If the total number of strands crosses $N_{max} = 1000$
   (a) Assign a probability $(1 - L_i^{-\gamma})$ to each strand, where $L_i$ is the length of i'th strand
   (b) Sample out $N_{max}$ number of strands from them according to those probabilities
6. Begin loop over time up to the duration of the dry phase
   (a) Calculate the total concatenation rates of the strands $\left(K_{con}^{Tot} = 4K_{con}N\right)$, where N is the number of strands
   (b) Draw a random-exponential time step dt based on this total rate
   (c) Add dt to the total time
   (d) If the total time after addition of dt is less than the duration of dry phase
      i. Choose a random strand
      ii. Extend it by adding a random monomer to it's 3' end
      iii. In parallel to this, draw a random number
      iv. If the random number is $< 16K_{con}dt$
         A. Create a dimer with two random monomers
         B. Add it to the population of strands
7. End loop over time

End loop over days.
